# The organizer of chromatin topology RIF1 ensures cellular resilience to DNA replication stress

**DOI:** 10.1101/669234

**Authors:** Cyril Ribeyre, Rana Lebdy, Julie Patouillard, Marion Larroque, Raghida Abou-Merhi, Christian Larroque, Angelos Constantinou

## Abstract

Eukaryotic genomes are duplicated from thousands of replication origins that fire sequentially forming a defined spatiotemporal pattern of replication clusters. The temporal order of DNA replication is determined by chromatin architecture and, more specifically, by chromatin contacts that are stabilized by RIF1. Here we show that RIF1 localizes in close proximity to newly synthesized DNA. In cells exposed to the DNA replication inhibitor aphidicolin, suppression of RIF1 markedly decreased the efficacy of protein isolation on nascent DNA (iPOND), suggesting that the iPOND procedure is biased by chromatin topology. RIF1 was required to limit the accumulation of DNA lesions induced by aphidicolin treatment and promoted the recruitment of cohesins in the vicinity of nascent DNA. Collectively, the data suggest that the stabilization of chromatin topology by RIF1 limits replication-associated genomic instability.

## Introduction

The duplication of a complete genome is a formidable task that must be perfectly controlled to avoid the transmission of mutations or chromosomal rearrangements to daughter cells. Two meters of DNA are packed and replicated in a human cell of about 10 μm of diameter. Hence, the spatiotemporal program of DNA replication is largely defined by the global organization of the nucleus (Marchal et al., 2019). DNA replication is initiated from defined regions of the genome called origins of replication. More than 30000 replication origins are required for the duplication of the human genome (Mechali, 2010). When replication forks stall, the firing of backup origins (also known as dormant origins) ensure the completion of DNA replication (Blow et al., 2011). The timing of replication is influenced by the 3D organization of chromatin architecture (Courbet et al., 2008; Foti et al., 2016; Klein et al., 2021). Cohesins influence origins firing locally (Guillou et al., 2010), yet without determining replication timing globally (Oldach and Nieduszynski, 2019), most likely via the formation of loops by extrusion (Davidson et al., 2019; Kim et al., 2019). RIF1, a conserved protein involved in telomeres capping, DNA double-strand break repair and chromatin organization, controls the timing of DNA replication (Cornacchia et al., 2012; Foti et al., 2016; Hayano et al., 2012; Klein et al., 2021; Mattarocci et al., 2016; Yamazaki et al., 2012). RIF1 determines replication timing via the stabilization of chromatin architecture (Foti et al., 2016; Kanoh et al., 2015; Klein et al., 2021; Yamazaki et al., 2013), and may regulate origin licensing owing to its interaction with PP1 phosphatase that would counteract DDK kinases (Dave et al., 2014; Hiraga et al., 2014; Mattarocci et al., 2014).

Throughout S phase, different nuclear patterns of replication foci reflect the orderly and sequential replication of chromatin domains (Chagin et al., 2016; Dimitrova and Berezney, 2002). Replication forks encounter a variety of impediments from both endogenous and exogenous sources (Lambert and Carr, 2013; Zeman and Cimprich, 2014). The slowing or stalling of replication forks by these impediments induces the activation of the checkpoint kinase ATR, which ensures that DNA synthesis within actively replicating chromosomal domains is completed before the duplication of a new chromosomal domain has started. ATR signaling delays the activation of late replication domains while promoting the firing of dormant origins within active replication domains (Blow et al., 2011). This suggests that the nuclear architecture contributes to cellular resilience to DNA replication stress. In support of this, Lamin A/C is required for the maintenance of chromosome integrity when the progression of replication forks is impeded by DNA lesions or upon nucleotide depletion (Singh et al., 2013). Furthermore, the association of Lamin A/C with the DNA polymerase clamp PCNA is critical for replication forks stability (Cobb et al., 2016). Hutchinson-Gilford progeria syndrome is caused by a mutation of the LMNA gene that leads to an aberrant Lamin A protein named progerin. The association of progerin with PCNA alters the nuclear distribution of PCNA, induces ATR activation and the formation of γH2A.X (Wheaton et al., 2017). In budding yeast, cohesins accumulate in the vicinity of replication forks upon treatment with hydroxyurea and are required for replication fork restart (Tittel-Elmer et al., 2012). These examples illustrate the links between replicative stress and nuclear structures, which remain incompletely understood.

The isolation of Proteins on Nascent DNA coupled with Mass Spectrometry (iPOND-MS) allows the identification of proteins localized in the vicinity of active replication forks (Aranda et al., 2014; Dungrawala et al., 2015; Lopez-Contreras et al., 2013; Lossaint et al., 2013; Sirbu et al., 2011; Sirbu et al., 2013). iPOND experiments performed under various experimental conditions have revealed components of the replication machinery (e.g. PCNA and DNA polymerases), proteins that accumulate near forks under stressful conditions (e.g. ATR and FANCD2), proteins that are required for the restoration of chromatin structures after passage of the replication fork (e.g. histones) and proteins that are playing a structural roles such as Lamin A (Alabert et al., 2014; Dungrawala et al., 2015; Lopez-Contreras et al., 2013; Lossaint et al., 2013; Ribeyre et al., 2016; Sirbu et al., 2011; Sirbu et al., 2013; Wheaton et al., 2017).

Here we provide evidence that during S phase, RIF1 is proximal to newly synthesized DNA. In cells exposed to the DNA polymerase inhibitor aphidicolin, RIF1 promotes the recruitment of the cohesin subunits SMC1 and SMC3 near replication forks and stabilizes replicating nucleoprotein clusters isolated by iPOND. We propose that the stabilization of chromatin architecture by RIF1 and cohesin limits the formation of DNA lesions caused by DNA replication impediments.

## Results

### iPOND coupled with mass spectrometry identifies proteins involved in nuclear organization

To identify new proteins in vicinity of replication forks we performed iPOND-MS using a highly sensitive last generation mass spectrometer (Sciex TripleTOF 5600+) and quantified the results using MaxQuant (Cox and Mann, 2008). We analyzed the data using Perseus (Tyanova et al., 2016) and took advantage of a volcano plot representation to visualize the proteins significantly enriched upon EdU pulse compared to a 2 hours thymidine chase (Figure 1A). As expected, most of the known proteins of the replisome (e.g., PCNA, RFC subunits, MCM1-6, FEN1 or DNA polymerases) were clearly enriched. We identified also a large number of proteins that were not previously described as replisome components that will be analyzed elsewhere. Interestingly, the iPOND-MS data revealed an enrichment of several cohesin subunits (SMC3, SMC1A, STAG2, RAD21, PDS5A and PDS5B) near forks (Figure 1A). Since cohesins are thought to play an architectural role at replication foci (Guillou et al., 2010), it is likely that they are not associated directly with individual replication forks but rather with chromatin domain undergoing replication. In contrast, Lamin B1 and Lamin B2 were not enriched after EdU pulse (Figure 1A), indicating that not all the structural components of the nucleus are localized in proximity of active replisomes. Interestingly, we identified RIF1 as a protein associated with nascent DNA (Figure 1A), consistent with previous studies (Alabert et al., 2014; Munden et al., 2018). We confirmed this data using an antibody directed against RIF1 (Figure 1B). This indicates that RIF1 localizes in the vicinity of active replication forks. Consistent with this, we detected RIF1 in immune-precipitates of the endogenous DNA polymerase clamp PCNA (Figure 1C).

**Figure 1:**
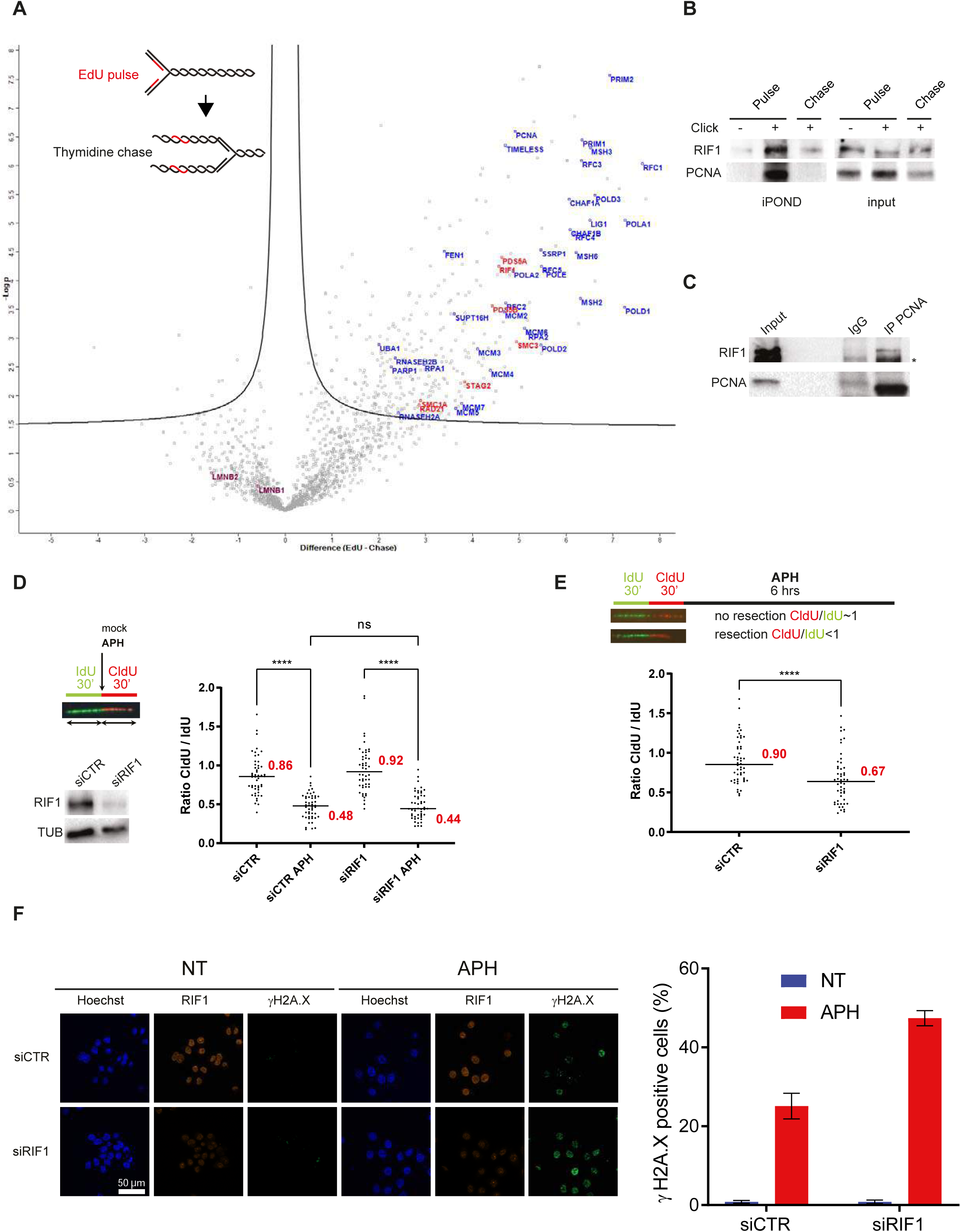
RIF1 is associated with nascent DNA and is required to limit DNA lesions in response to prolonged aphidicolin treatment. **A**. iPOND coupled with mass spectrometry. HeLa S3 were pulse-labeled with EdU or pulse-labelled with EdU then subjected to a 120 min thymidine chase the subjected to iPOND and analyzed by mass-spectrometry. Label free quantification was performed using MaxQuant (Cox and Mann, 2008) and statistical analysis using Perseus (Tyanova et al., 2016). Pulse experiments have been performed 6 times and chase experiments 4 times. Examples of replisome-specific proteins are indicated on the right side of the figure above the line. Full proteins list is available in Supplementary Table 1. **B**. Indicated proteins were isolated by iPOND and detected by Western blotting. HeLa S3 cells were pulse-labelled with EdU for 15 min and chased with thymidine for 120 min. In no click biotin-TEG azide was replaced by DMSO. **C**. Western-blot analysis of indicated proteins after immunoprecipitation with an antibody directed against PCNA or against mouse IgG. **D**. DNA fibers labelling and Western-blot analysis of RIF1 depletion. HeLa S3 cells were labelled for 30 min with IdU and then for 30 min with CldU in the absence or presence of 0.05 μM aphidicolin (APH) in the cell culture medium. Graphic representation of the ratios of CldU versus IdU tract length. For statistical analysis Mann-Whitney test was used; ns, non-significant, ****p<0.0001. The horizontal bar represents the median with the value indicated in red. 50 replication tracts were measured for each experimental condition. **E**. Analysis of DNA resection using DNA fibers labelling. HeLa S3 cells were labelled for 30 min with IdU and then for 30 min with CldU. 1 μM aphidicolin (APH) was added in the cell culture medium for 6 hours. Graphic representation of the ratios of CldU versus IdU tract length. For statistical analysis Mann-Whitney test was used; ****p<0.0001. The horizontal bar represents the median with the value indicated in red. 50 replication tracts were measured for each experimental condition. **F**. Immunofluorescence analysis of γH2A.X and RIF1 in HeLa S3 cells with siRNA against control or RIF1 in presence or absence of aphidicolin (APH) for 24 hours. Graphic representation of the percentage of γH2A.X positive cells based on 3 independent experiments.

### RIF1 protects the integrity of replication forks upon prolonged replicative stress

Although RIF1 located near active replisomes, suppression of RIF1 did not alter significantly the progression of replication forks (SupFig1A), consistent with previous studies (Cornacchia et al., 2012; Ray Chaudhuri et al., 2016). A higher frequency of stalled forks, however, is observed in *rif1*^-/-^ DT40 cells (Xu et al., 2010), suggesting that RIF1 could be important for fork progression in some contexts. Consistent with this, several studies have detected the activation of the checkpoint effector kinase Chk1 in RIF1-depleted cells (Chapman et al., 2013; Foti et al., 2016). We confirmed that Chk1 was active by phosphorylation on Serine 345 upon suppression of RIF1 by means of siRNAs (SupFig1B). We observed also that RPA32 was phosphorylated on Ser4/8, suggesting that RIF1 depleted cells accumulate DNA lesions (SupFig1B). Interestingly, RIF1 recruitment at replication forks is slightly increased upon hydroxyurea (HU) treatment to limit DNA2-mediated DNA resection and DNA lesions (Garzón et al., 2019; Mukherjee et al., 2019; Ray Chaudhuri et al., 2016). Consistent with this, DNA lesions, genetic instability and HU sensitivity are increased upon RIF1 impairment (Buonomo et al., 2009; Mukherjee et al., 2019; Xu et al., 2010). This raises the possibility that the stabilization of chromatin topology by RIF1 limits replication-associated DNA lesions under stressful conditions. To test this, we analyzed if RIF1 loss had any impact on replication fork dynamics in the presence of aphidicolin (APH). We labelled cells for 30 minutes with IdU and then for 30 minutes with CldU in presence of a low dose (0.05 μM) of APH. As expected, the ratio of the lengths of CldU versus IdU tracts was close to 1 in control conditions and reduced by half in presence of APH (Figure 1D, SupFig1C). The status of RIF1 did not change the ratios of CldU/IdU tracts (Figure 1D, SupFig1C) indicating that RIF1 depletion does not play any major role in early responses to APH. As RIF1 is protecting HU-stalled forks from nucleases degradation (Garzón et al., 2019; Mukherjee et al., 2019), we tested if this was also the case when replication forks were blocked with APH. To do so, we treated cells 6 hours with a high dose (1 μM) of APH after 30 min sequential labelling of IdU and CldU and measured the ratio between the lengths of CldU and IdU tracts. The ratio was close to 1 in cells treated with a control siRNA, and below 1 in RIF1 depleted cells, confirming that RIF1 is indeed protecting APH-stalled forks (Figure 1E). Consistent with this, prolonged treatment (24 hours) with APH, increased the percentage of γ-H2A.X-positive cells to almost 2-fold (Figure 1E) and decreased by two-fold the ability of replication forks to restart (SupFig1D). Altogether, these data indicate that RIF1 limits the formation of DNA lesions under stressful conditions.

### RIF1-dependent loss of replication organization induces DNA lesions

Despite its role in protection of stalled replication forks (see above), RIF1 recruitment at forks does not increase significantly in response to HU (Mukherjee et al., 2019) compared to proteins such as ATR, 9-1-1, TopBP1 or FANCD2/FANCI (Dungrawala et al., 2015; Lossaint et al., 2013). Therefore, we hypothesize that the impact of RIF1 on nascent DNA protection may not reflect a direct role at stalled replication forks. This is supported by several articles showing that RIF1 is crucial for the organization of higher-order chromatin domains and for the establishment of the replication timing program (Foti et al., 2016; Klein et al., 2021; Moriyama et al., 2018; Yamazaki et al., 2012). Remarkably, the mid-S pattern is selectively loss upon RIF1 impairment (Yamazaki et al., 2012), this effect was attributed to the impact of RIF1 in replication timing. However, we noticed that these experiments have been performed in cells synchronized with thymidine block and released into S-phase. It is well established that synchronization with thymidine block perturbs the pool of nucleotides and induces DNA damage (Kurose et al., 2006). Thus, we hypothesized that the absence of mid-S pattern in RIF1-depleted cells synchronized using a thymidine block could reflect a defect in the maintenance of chromatin topology during DNA replicative stress. To test this, we compared the frequency of each pattern in asynchronous conditions and in cells synchronized with thymidine block and released into S-phase upon RIF1 depletion (Figure 1A). In synchronous condition, we were able to reproduce the results of Yamazaki et al. and observed the disappearance of the mid-S pattern upon RIF1 depletion (Figure 2B, 2C). Surprisingly, in asynchronous conditions, we found that RIF1 depletion did not alter the occurrence of the mid-S pattern (Figure 2B, 2C). Importantly, and as already observed (Yamazaki et al., 2012), cell-cycle distribution was not significantly affected in absence of RIF1 in synchronous or asynchronous conditions (SupFig2A). This result suggests that the disappearance of the mid-S pattern in RIF1 depleted cells is a consequence of the synchronization procedure. This observation cannot be solely explained by the difference in replication timing since it should be also observed in asynchronous cells. To test if synchronization procedure increases the level or replicative stress, we analyzed the level of the marker of DNA damage γ-H2A.X. In an asynchronous population of cells, the depletion of RIF1 had no impact on the percentage of γ-H2A.X positive cells (Figure 2B, 2D). As expected, the percentage of γ-H2A.X positive cells increased 2 hours after release from the thymidine block. Strikingly, inactivation of RIF1 tripled the percentage of γ-H2A.X positive cells in the same conditions (6.9% in control versus 24.1% in shRIF1 (1) and 19.1% in shRIF1 (2)). We conclude that the disappearance of the mid-S pattern upon RIF1 depletion correlates with the formation of DNA lesions. The thymidine block procedure is affecting the pool of dNTPs and therefore should have a direct impact on the progression of replication forks that might be exacerbated in the absence of RIF1. To test this, we monitored the phosphorylation of Chk1 on Serine 345. In control condition, we observed a mild phosphorylation of Chk1 on Serine 345, in line with the higher level of γ-H2A.X (Figure 2E). Interestingly, we observed a strong level of Chk1 phosphorylation in RIF1-depleted cells 2 hours after release from the thymidine block (Figure 2E). In asynchronous conditions, suppression of RIF1 did not significantly alter the progression of replication forks (SupFig1A). Two hours after release from a thymidine block, however, replication tracts were longer in the absence of RIF1 (Figure 2F, SupFig2B). Unrestrained DNA synthesis would yield single-stranded DNA gaps detected and signaled by ATR, consistent with higher level of Chk1 and H2A.X phosphorylation. We propose that the occurrence of DNA lesions during prolonged replicative stress observed in RIF1 depleted cells is a consequence of alterations in the organization of replicated chromatin domains.

**Figure 2:**
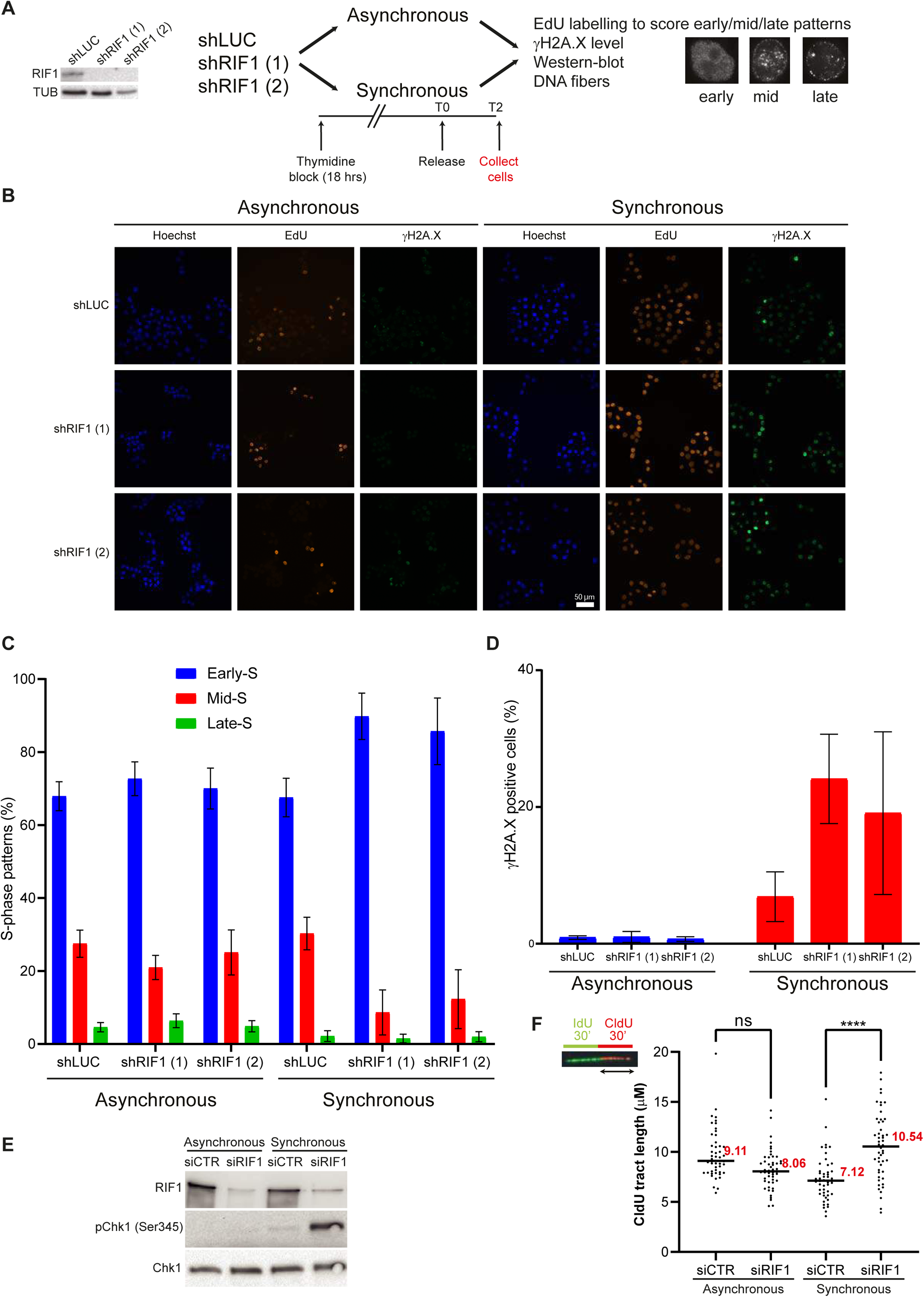
RIF1 depletion alters S-phase organization and yields DNA lesions. **A**. Experimental setup to study the impact of synchronization procedure in HeLa S3 cells depleted or not for RIF1. The efficacy of RIF1 depletion using 2 different shRNA is shown. For synchronization, cells were grown 18 hours in presence of 2mM thymidine then released into S-phase for 2 hours. Cells were then subjected to immunofluorescence, western-blot or DNA fibers analysis. **B**. Immunofluorescence analysis of γH2A.X and EdU in asynchronous and synchronous HeLa S3 cells expressing shRNAs against luciferase or RIF1. **C**. Graphic representation of the frequency of replication patterns (Late-S, Mid-S and Early-S) based on at least three independent experiments for each condition. **D**. Quantification of γH2A.X intensity within nucleus stained with Hoechst using CellProfiler based on at least three independent experiments for each condition. **E**. Western-blot analysis of Chk1 phosphorylation on Serine 345 upon RIF1 depletion **E**. DNA fibers assay. HeLa S3 cells were labelled for 30 min with IdU and then for 30 min with CldU. Graphic representation of CldU tracts lengths. For statistical analysis Mann-Whitney test was used; ns, non-significant, ****p<0.0001. The horizontal bar represents the median with the value indicated in red. At least 50 replication tracts were measured for each experimental condition.

### RIF1 impairment reduces iPOND efficiency in presence of replicative stress

We showed that prolonged treatment with APH or thymidine yields high level of γH2A.X in RIF1-depleted cells. Importantly, after a thymidine block, the increase of γH2A.X signal correlates with alterations of DNA replication patterns. Since APH has also been widely used for cell synchronization, it is highly probable that the increase level of DNA lesions is absence of RIF1 in APH treated cells is also due to a defect in the maintenance of chromatin topology. Alternatively, the data could reflect a role for RIF1 in G1 cells rather than in S phase. To understand this in more details, we performed a series of experiments in RIF1-depleted cells using short treatments with aphidicolin (Figure 3A). First, we performed iPOND assay and probed isolated proteins by western-blotting (Figure 3B). Under standard cell culture conditions, the efficacy of PCNA isolation with nascent DNA in RIF1-depleted cells was similar to that of control cells (Figure 3B). As expected, a 30 min treatment with a low dose of APH (0.1 μM) induced the recruitment of BRCA1 and TopBP1 on nascent DNA (Figure 3B). Strikingly, in RIF1-depleted cells treated with APH, the efficacy of PCNA recovery (as well as BRCA1 and TopBP1) with nascent DNA diminished dramatically (Figure 3B), an observation we could reproduce with PCNA and other replisomes components (SupFig3A). This observation could be the consequence of a massive decrease in EdU incorporation that would impair proteins recovery. To test this hypothesis, we measured EdU incorporation using immunofluorescence. As expected APH treatment reduces strongly EdU incorporation but this effect was similar in RIF1-depleted cells, suggesting that EdU incorporation is not impaired (Figure 3C). In addition, and consistent with Figure 1D, DNA fibers experiment performed in the exact same cell line than the one used for iPOND indicate that DNA synthesis is occurring, although at lower pace, in response of APH independently of the presence of RIF1 (Figure 3D). Thus, a defect in DNA synthesis does not account for the reduced isolation of EdU-bound proteins from RIF1-depleted cells. Furthermore, we detected similar levels of the replisome-associated proteins MSH2 and MCM7 in PCNA immune-precipitates from control and RIF1-depleted cells (SupFig3B), suggesting that RIF1 is not required for replisome stability and replication fork progression. As this step, the most reasonable hypothesis is that in presence of replicative stress, suppression of RIF1 reduces the efficacy of capture of EdU-associated proteins. To generalize this observation, we performed the exact same iPOND experiment than in Figure 3B but analyzed the pulldowns using mass spectrometry (Figure 3E). In comparison with control cells, the treatment of RIF1-depleted cell with APH markedly reduced the abundance of the replication factors PCNA, MSH6, DPOD1, FEN1 and RFC4 captured by iPOND (Figure 3E). By contrast, changes in the efficacy of streptavidin pulldowns were not observed for mitochondrial proteins such as NDUS1, NDUS3, P5CR2 and SDHA, which are also isolated by iPOND (SupFig3C). To generalize this observation to the whole replisome, we summed the peptides intensities of all replisome proteins listed in a previous study (Lopez-Contreras et al., 2013). In control cells (shLUC), APH treatment moderately affected the recovery of replisome components (SupFig3D). By contrast, APH had a severe impact on the recovery of replisome components from RIF1-depleted cells (∼50% decrease for shRIF1 (1) and ∼33% decrease for shRIF1 (2)). The data indicate that APH treatment reduces the probability to capture proteins associated with EdU-labelled DNA in RIF1-depleted cells.

**Figure 3:**
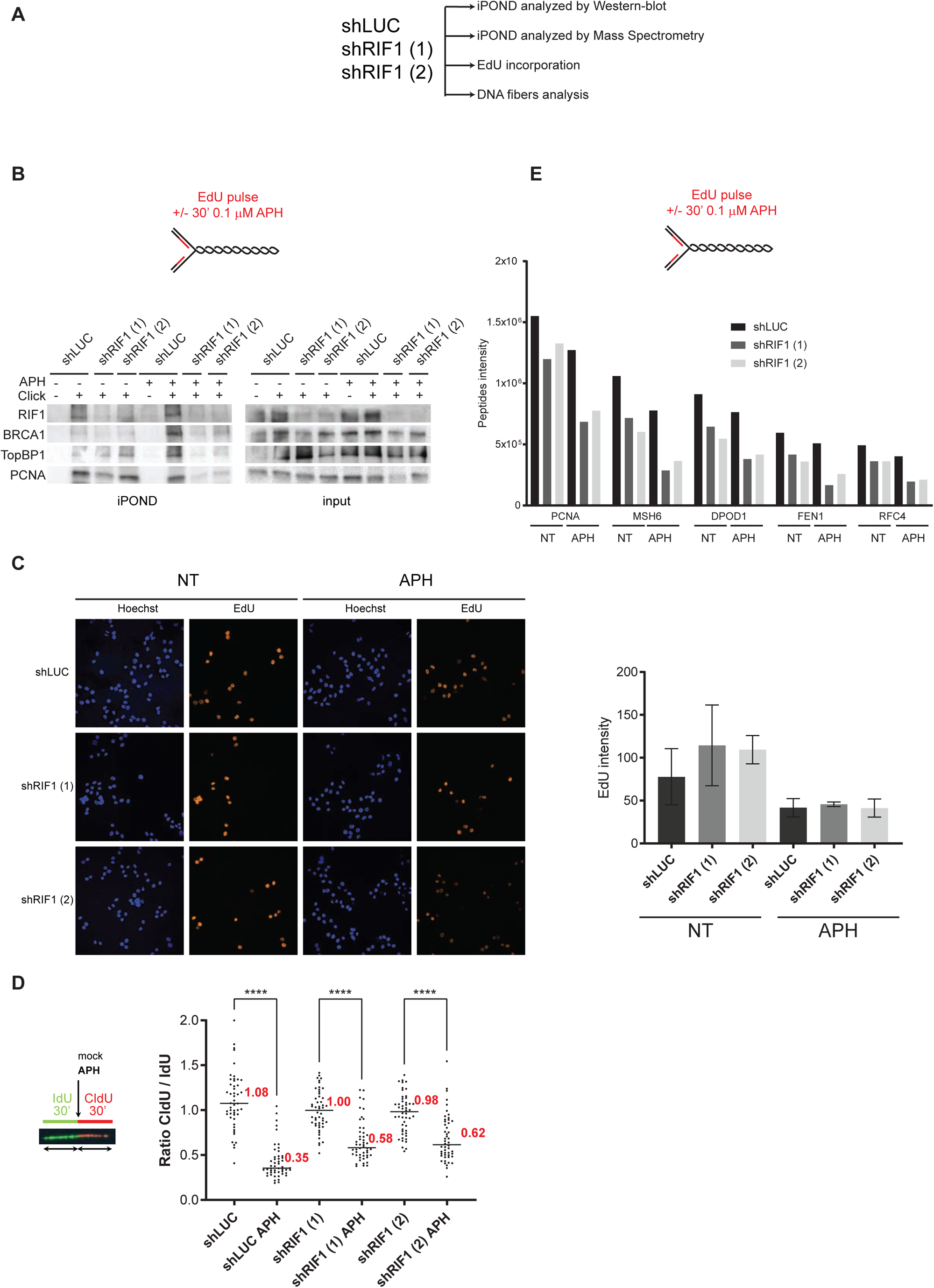
RIF1 loss reduces the efficacy of proteins isolation on nascent DNA. **A**. Experimental set-up. **B**. iPOND experiment. HeLa S3 cells (with shLUC or two different shRIF1) were labelled with EdU for 15 min or for 30 min with 0.1 μM aphidicolin (APH). Indicated proteins were analyzed by western-blotting. In no click samples biotin-TEG azide was replaced by DMSO. **C**. Analysis of EdU incorporation using microscopy in HeLa S3 cells with shRNA against luciferase or RIF1. EdU was incorporated in cells during 15 min with or without 0.1 μM aphidicolin (APH). Quantification of EdU intensity within nucleus stained with Hoechst was performed using CellProfiler and is represented on the histogram. Error-bars corresponds to the average values of three independent experiments. **D**. DNA fibers labelling. HeLa S3 cells were labelled for 30 min with IdU and then for 30 min with CldU in the absence or presence of 0.05 μM aphidicolin (APH) in the cell culture medium. Graphic representation of the ratios of CldU versus IdU tract length. For statistical analysis Mann-Whitney test was used; ****p<0.0001. The horizontal bar represents the median with the value indicated in red. At least 50 replication tracts were measured for each experimental condition. **E**. iPOND-MS experiment. HeLa S3 cells (with shLUC or two different shRIF1) were labelled with EdU for 15 min or for 30 min EdU with 0.1 μM aphidicolin (APH). Quantification of peptides intensity corresponding to the indicated proteins is represented.

### The efficacy of iPOND is biased by chromatin topology

How can we explain that in cells exposed to aphidicolin, RIF1 depletion decreases the efficacy of iPOND without affecting EdU incorporation? To answer to this question, one has to take into consideration that the association of proteins such as cohesins or RIF1 with EdU may be indirect and determined by chromatin topology. Consistent with this, methods that are using formaldehyde crosslinking such as ChIP or chromosome conformation capture are indeed dependent on nuclear organization. To test if iPOND efficiency is biased by chromatin organization, we took advantage of the distinct and characteristic patterns formed by replicons labelled with EdU (Dimitrova and Berezney, 2002). In early S-phase (replication of euchromatin), the EdU pattern is poorly clustered. Clusterization then increases in mid-S phase (replication of facultative heterochromatin) and is even stronger in late S-phase (replication of constitutive heterochromatin). We synchronized HeLa S3 cells using a simple thymidine block procedure and released the cells in fresh media without thymidine (Figure 4A). We added EdU for 15 minutes just before release (T0) and then 2 hours (T2), 4 hours (T4) and 8 hours (T8) after release (Figure 4A). We verified the synchronization procedure by flow cytometry using double labelling with EdU and propidium iodide (Figure 4B). As expected at T0 the majority (∼80%) of the cells were in G1. Two and four hours after release (T2 and T4) most of the cells (∼80%) were in S-phase. After 8 hours (T8), cells entered G2 and the number of S-phase cells decreased (∼25%). We then performed iPOND experiment on synchronized and non-synchronized cells. At T0, the PCNA signal was barely detectable, as expected, and comparable to the control (minus click) of the asynchronous conditions (Figure 4C). By contrast, we could observe a clear PCNA signal after the EdU-Biotin click reaction in non-synchronized condition. At T2 and T4 the PCNA signal became detectable. Surprisingly the strongest signal was observed at T8 despite the fact that the number of cells in S-phase is lower than in T4 and T2 (Figure 4C). This observation is also true for MCM7 and H3 (Figure 4C) and is reproducible (Figure 4D). This result indicates that the efficacy of protein isolation on nascent DNA does not correlate directly with the number of cells in S-phase. Therefore, we propose that the recovery of replisomes components by iPOND is strongly dependent on the organization of replicated chromatin domains (Sup Fig4). Thus, we suggest that the reduced efficacy of EdU-biotin streptavidin pulldowns in the iPOND procedure and the accumulation of markers of DNA damage in RIF1 depleted cells exposed to aphidicolin reflects a role for RIF1 in the stabilization of chromatin topology under DNA replication stress conditions.

**Figure 4:**
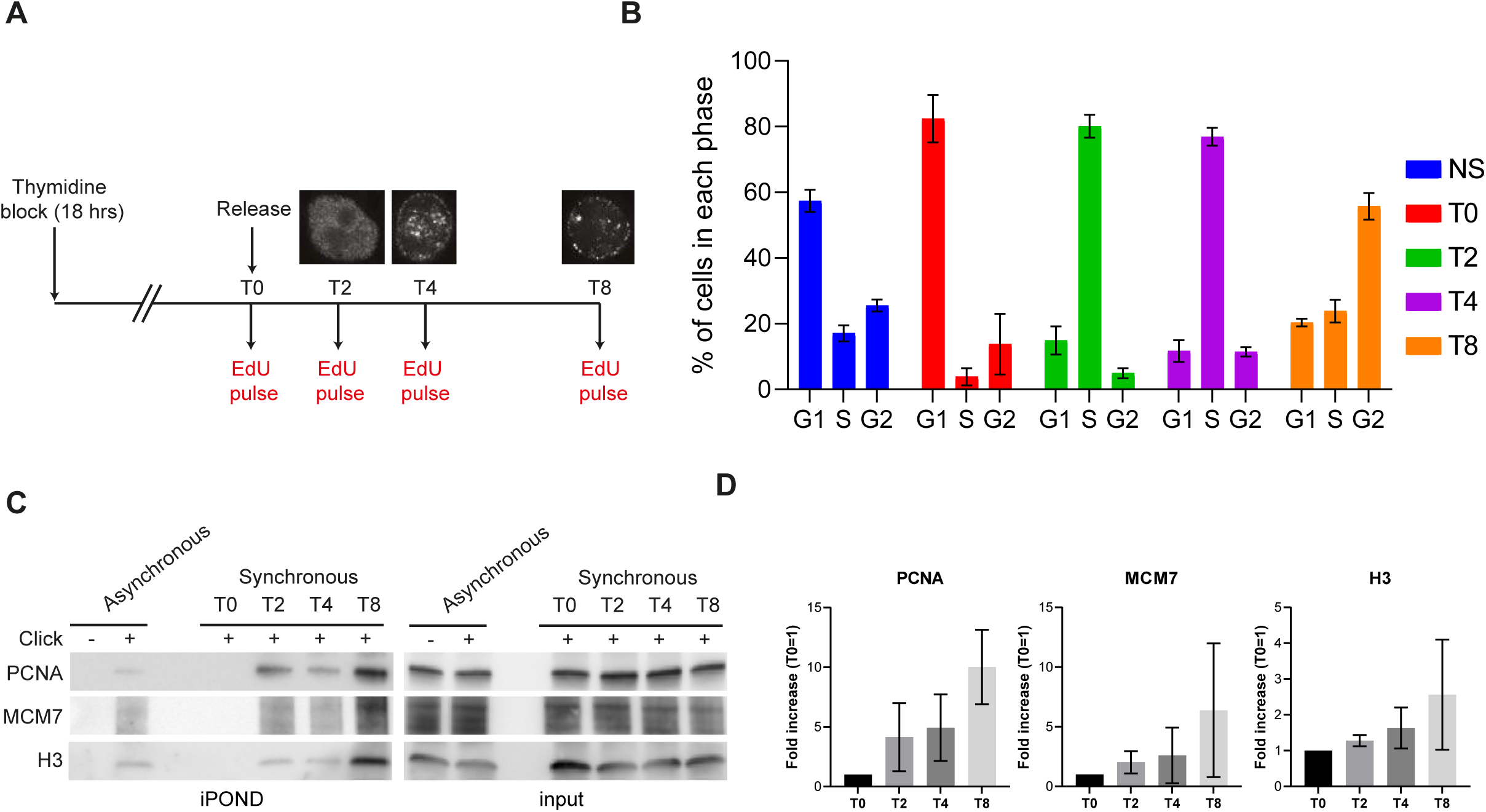
iPOND proteins recovery is biased by replication organization. **A**. Experimental set-up. HeLa S3 cells were submitted to thymidine block during 18 hours and released into S-phase. Cells were collected at T0 (G1), T2 (Early-S), T4 (Mid-S) and T8 (Late-S) after 15 min EdU pulse for iPOND and flow cytometry. Replication patterns showing the different phases are represented. **B**. The percentage of cells in each phase was analyzed using flow cytometry. The error-bars represent the variations within 3 independent experiments. **C**. iPOND experiment performed on unsynchronized and synchronized cells and analyzed by western-blot using antibodies directed against the indicated proteins. In no click samples, biotin-TEG azide was replaced by DMSO. **D**. Quantification of the indicated proteins in iPOND based on at least 3 independent experiments, T0 was used for normalization.

### RIF1 depletion impairs the loading of SMC1 and SMC3 at forks in presence of replicative stress

RIF1 stabilizes chromatin topology via its intrinsic capacity to bridge molecules (Mattarocci et al., 2017) and may promote the recruitment of additional proteins involved in the organization of chromatin topology such as the cohesin complex. Indeed, cohesin are associated with replication forks in basal conditions (Figure 1A) and in response to replicative stress (Ribeyre et al., 2016; Tittel-Elmer et al., 2012). In addition, cohesins cooperate with RIF1 in the stabilization of chromatin topology at sites of DNA double-strand breaks (Ochs et al., 2019), and organize DNA repair foci via a mechanism of loop extrusion at both sites of the DNA breaks (Arnould et al., 2021). Since RIF1 depletion diminishes the efficacy of the iPOND procedure, we used, as an alternative method, a proximity ligation assay (PLA, Figure 5A) to analyze the loading of cohesins subunits in the vicinity of nascent DNA (Petruk et al., 2017; Petruk et al., 2012; Roy et al., 2018). We first validated the method using PCNA as positive control. As expected, we detected PCNA-EdU proximity signals in cells after the coupling of EdU and biotin, specifically (SupFig5A). We then analyzed the recruitment of SMC1 to nascent DNA in presence of 0.1μM APH. In control conditions, we observed a clear PLA signal between EdU and SMC1 confirming that SMC1 is recruited near stalled replication forks (Figure 5B). Interestingly, the signal of proximity between EdU and SMC1 was reduced in RIF1-depleted cells (Figure 5B, 5C, SupFig 5B). Consistent with this, the localization of SMC3 to stalled forks was also dependent on RIF1 (SupFig5C). By contrast, RIF1 suppression had no impact on EdU-PCNA proximity signal (Figure 5B, 5C, SupFig 5B). We verified that the suppression of RIF1 did not reduce the level of SMC3 and SMC1 expression (SupFig5D). Thus, we conclude that RIF1 contributes to the loading of the cohesins subunits SMC1 and SMC3 near stalled DNA replication forks.

**Figure 5:**
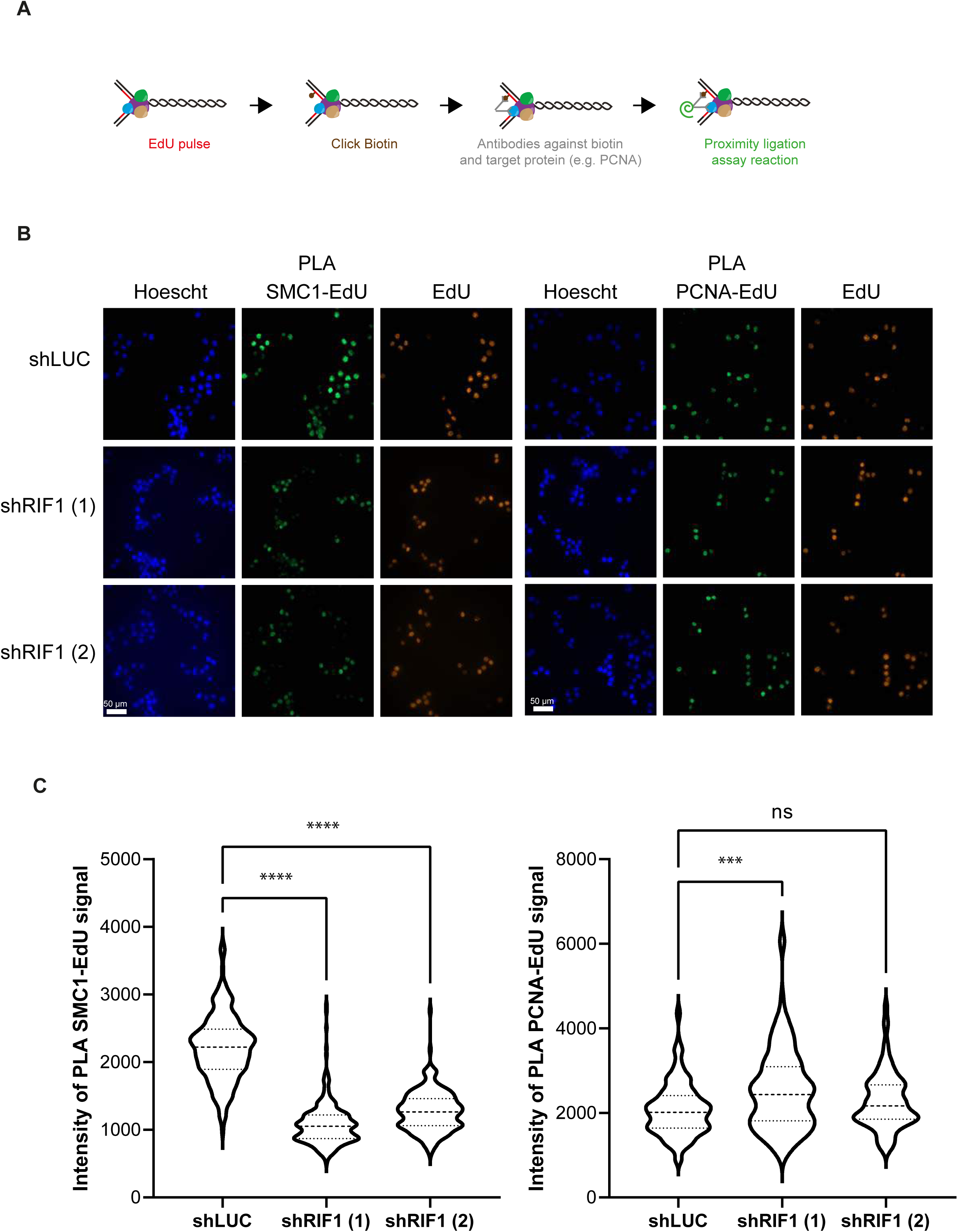
RIF1 is required for full recruitment of cohesin subunits at stalled forks. **A**. Scheme explaining the principle of proximity ligation assay (PLA) between EdU and replisome components. **B**. Immunofluorescence analysis of PLA signal between EdU and SMC1 and between EdU and PCNA upon 30 min treatment with 0.1 μM APH in HeLa S3 cells expressing shRNAs against luciferase or RIF1. EdU-positive cells were labelled with Alexa-Fluor 555. **C**. The level of PLA signal within the nucleus was quantified using CellProfiler. Graphical representation of the PLA signal, at least 100 cells were quantified in each condition. For statistical analysis Mann-Whitney test was used; ns, non-significant, ****p<0.0001; ***p<0.001.

## Discussion

RIF1 was originally discovered more than 25 years ago in budding yeast as a negative regulator of telomere elongation (Hardy et al., 1992). It is now clearly established that RIF1 is a highly conserved protein (Sreesankar et al., 2012) involved in telomeres protection, DNA replication, DNA double-strand break repair, transcription and heterochromatin formation (Mattarocci et al., 2016). The links between the seemingly disparate functions of RIF1 may stem for the function of RIF1 in the stabilization of chromatin topology (Arnould et al., 2021; Klein et al., 2021). Here we provide evidence that the organization of chromatin architecture by RIF1 ensures chromosome stability during DNA replication stress. This model is based on the following findings: (1) RIF1 locates near replication sites in basal conditions (2) DNA replication stress in RIF1-depleted cells modifies S phase patterns and increases the level of the DNA damage marker γH2A.X (3) Suppression of RIF1 strongly affects the organization of DNA replication in response to replicative stress. (4) RIF1 may exert this function in coordination with the cohesin complex. Our model is consistent with the finding that RIF1 bridges proximal DNA molecules (Mattarocci et al., 2017) and creates a protective structure around DBSs (Ochs et al., 2019). Thus, we propose that the chromatin organizing function of RIF1 ensures DNA replication under stressful conditions (Figure 6). By analogy with its function at yeast telomeres, we would like to propose that RIF1 is protecting replication domains (Figure 6).

**Figure 6:**
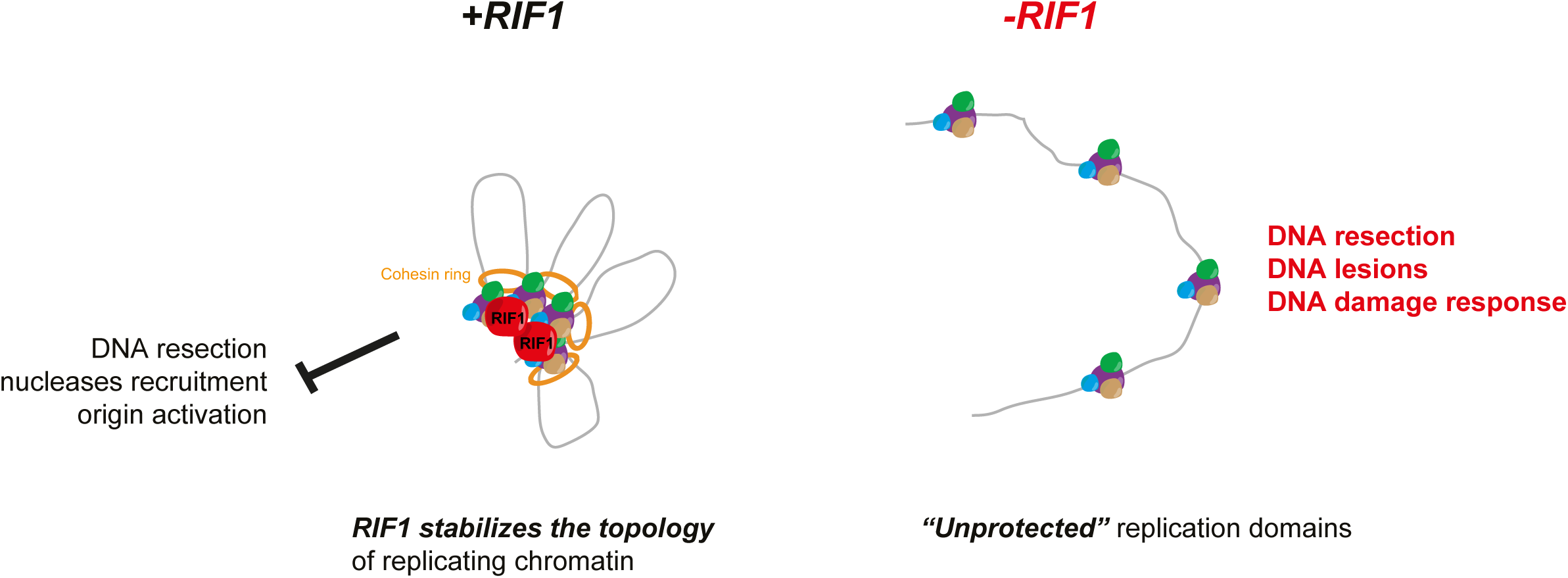
Model to explain the role of RIF1 in the organization of replication factories. RIF1 stabilizes chromatin topology during DNA replication thus preventing DNA resection by nucleases or excessive origins activation. This could direct, thanks to its capacity to interact with DNA, or/and via the recruitment of cohesin ring. In its absence, the replication domains are unprotected leading to DNA resection, DNA lesions and activation of DNA damage response.

The association of RIF1 with the replication forks has been previously observed by other groups (Alabert et al., 2014; Her et al., 2018; Munden et al., 2018). We confirmed that suppression of RIF1 has no measurable effect on replication forks progression under standard conditions or in response to short treatment with replicative stress (Cornacchia et al., 2012; Ray Chaudhuri et al., 2016; Xu et al., 2010) despite the fact that RIF1 loss induces Chk1 phosphorylation on Ser345 (Chapman et al., 2013; Foti et al., 2016). Interestingly, we found that two hours after release from a thymidine block, replication tracts are longer in the absence of RIF1 and phosphorylation levels of Chk1 on Ser345 and H2A.X on Ser139 are increased. One possibility is that in the absence of RIF1, the disorganization of chromatin domains during DNA replication results in the accumulation of single-stranded DNA. The accumulation of DNA lesions likely underpins the increased sensitivity of RIF1 defective cells to inhibitors of DNA replication (Buonomo et al., 2009; Feng et al., 2013; Xu et al., 2010; Xu et al., 2017).

RIF1 is recruited by 53BP1 at DSBs to prevent homologous recombination and favor NHEJ (Chapman et al., 2013; Di Virgilio et al., 2013; Escribano-Diaz et al., 2013; Zimmermann et al., 2013). Based on this, it has been proposed that RIF1 could be recruited by 53BP1 to protect stalled forks independently of BRCA1 (Xu et al., 2017). These data are raising the possibility that 53BP1 contributes to the recruitment of RIF1 at replication forks in basic conditions and in response to replicative stress. However, RIF1 recruitment is not impacted by 53BP1 depletion (Her et al., 2018) and RIF1, but not 53BP1, protects nascent DNA from degradation (Ray Chaudhuri et al., 2016) suggesting that the presence of RIF1 at replication forks is independent of 53BP1, consistent with it capacity to form higher order structures in budding yeast (Mattarocci et al., 2017).

Our model is consistent with the observation that RIF1 protects stalled replication forks from resection by nucleases, perhaps via the creation of a compartment that prevent their recruitment (Garzón et al., 2019; Mukherjee et al., 2019; Ray Chaudhuri et al., 2016). A role for RIF1 in safeguarding the stability of replicated domains may also explain how RIF1 controls the activation of dormant origins in response to replicative stress (Hiraga et al., 2017) and prevents the formation of anaphase bridges (Hengeveld et al., 2015 ; Zaaijer et al., 2016). RIF1 depletion has a strong impact on replication timing (Cornacchia et al., 2012; Foti et al., 2016; Yamazaki et al., 2012). The action of RIF1 on the replication timing program may result from the regulation of DDK kinase activation through RIF1 interaction with the PP1 phosphatase (Dave et al., 2014; Hiraga et al., 2014; Mattarocci et al., 2014) or through its ability to bind G-quadruplexes and to organize chromatin topology (Kanoh et al., 2015). Since the loss of RIF1 induces drastic changes in nuclear organization revealed by chromosome conformation capture methods (Foti et al., 2016), we favor the hypothesis that the impact of RIF1 on replication timing is a consequence of impaired nuclear organization rather than of a defect in the control of DDK kinases. The later hypothesis is supported by recent evidence based on Hi-C chromosome conformation capture experiments showing that RIF1 is necessary to enforce chromosome interaction hubs that determine the replication-timing program (Klein et al., 2021). This model could explain why suppression of RIF1 perturbs transcription and heterochromatin formation (Dan et al., 2014; Hiraga et al., 2017; Klein et al., 2021). Since the recruitment of cohesins at stalled forks is dependent on RIF1, it is tempting to speculate that RIF1 might ensures the stabilization of replicating chromatin domains in coordination with cohesin. Consistent with this, the depletion of cohesin subunits mimics topological alterations at DSBs caused by the depletion of RIF1 (Ochs et al., 2019). In contrast, induced degradation of SCC1 did not impact the patterns of replication (Oldach and Nieduszynski, 2019). We favor a model where the cohesin complex is recruited by RIF1 directly at stalled forks to maintain the local organization of chromatin, with no impact on the general organization of DNA replication (Ribeyre et al., 2016; Tittel-Elmer et al., 2012).

Finally, this study illustrates an as yet unforeseen application of iPOND (or iPOND-related methods based on formaldehyde crosslinking). It is generally assumed that the iPOND method captures proteins associated with individual replisomes distributed along a linear DNA template. Here we show that the iPOND method is not only efficient to isolate replisome components but also to capture structural components of replicating chromatin domains stabilized by formaldehyde crosslinking. Future studies using iPOND and other method should provide new insights into the role of the nuclear organization in DNA replication.

## Acknowledgements

We thank all the lab members for comments and suggestions on the project and on the manuscript. We are grateful to Antoine Aze for critical reading of the manuscript. We thank Marie-Pierre Blanchard and the MRI microscopy platform for their support. We acknowledge the support of the Site de Recherche Intégrée sur le Cancer Montpellier Cancer grant INCa_INSERM_DGOS_12553. This work was supported by a Jeunes Chercheuses Jeunes Chercheurs (JCJC) grant (REPLIBLOCK ANR-17-CE12-0034-01) from the Agence National de la Recherche (ANR) to Cyril Ribeyre and by grants from la Fondation ARC pour la Recherche sur le Cancer (PGA1 RF20180206787) and Merck Sharp and Dohme Avenir (GnoSTic) to Angelos Constantinou. Rana Lebdy is funded by fellowships from Azm & Saade Association and Fondation ARC pour la recherche sur le cancer.

## Author Contributions

Cyril Ribeyre: conceptualization, data curation, supervision, formal analysis, funding acquisition, investigation, visualization, methodology, project administration and writing original draft, review and editing.

Rana Lebdy: conceptualization, data curation, formal analysis, investigation, visualization, methodology and draft review and editing

Julie Patouillard: formal analysis, investigation, visualization and methodology.

Marion Larroque: data curation, formal analysis, investigation and methodology.

Christian Larroque: data curation, formal analysis and methodology.

Raghida Abou-Merhi: supervision, project administration.

Angelos Constantinou: conceptualization, data curation, supervision, funding acquisition, visualization, methodology, project administration and writing original draft, review and editing.

## Methods

### Cell lines

HeLa S3 (obtained from ATCC) cells were cultured in Dulbecco’s modified Eagle’s media (DMEM). Culture media was supplemented with 10% fetal bovine serum (Biowest) and penicillin/streptomycin (Sigma-Aldrich). Cells were incubated in a 5% CO_2_ at 37°C. For thymidine block experiments cells were treated18 hours with 2 mM thymidine, washed then release into normal media.

### Gene silencing

For RIF1 depletion siRNA oligonucleotides were purchased from Dharmacon (M-027983-01-0005) and transfected using INTERFERin (Polypus transfection). Anti-RIF1 shRNAs (1) and (2) and anti-luciferase shRNA were cloned in pSUPER-EBV and transfected using Lifofectamine 2000 (Thermo-Fisher). Stable cell lines were selected using puromycin.

### Western-blot

The proteins were resolved by SDS-PAGE using home-made or precast gels (Bio-Rad) and transferred to a nitrocellulose membrane (GE Healthcare or Bio-Rad). Antibodies against the following proteins were used: Ser345 Phospho-Chk1 (Cell Signaling Technology 2348), Chk1 (Santa Cruz sc-8408), PCNA (Sigma-Aldrich P8825), Ser4/8 Phospho-RPA32 (A300-245A), RPA32 (Calbiochem NA18), TopBP1 (Bethyl A300-111A), histone H3 (Abcam ab62642) BRCA1 (Santacruz sc-642), RIF1 (Bethyl A300-568A-M) and MCM7 (Abcam ab2360).

### Co-Immunoprecipitation

Cells were incubated for 30 min in ice in high salt buffer (50 mM Tris Ph 7.5, 300 mM NaCl, 1% Triton, 1 mM DTT). After 10 min centrifugation at 14000g, supernatant was incubated with anti PCNA antibody (Sigma-Aldrich, P8825) or IgG Rabbit (Calbiochem NI01) overnight at 4°C. Magnetic beads coupled with protein G (Life 10004D) were added for 1 hour and washed 5 times with washing buffer (10 mM Hepes, 100 mM KOAc, 0.1 mM MgOAc). Beads were boiled in Laemmli buffer and supernatants were analyzed by Western-blot.

### Isolation of proteins on Nascent DNA (iPOND)

iPOND was performed largely as described in (Lossaint et al., 2013; Ribeyre et al., 2016). Briefly, HeLa S3 cells were pulse labeled with 10 μM EdU for 5-15 min and a 120 min chase was performed with 10 μM thymidine. Cells were fixed with 1% formaldehyde for 5 min followed or not by quenching of formaldehyde by 5 min incubation with 0.125 M glycine. Fixed samples were collected by centrifugation at 2000 rpm for 3 min, washed three times with PBS and stored at -80°C. Cells were permeabilized with 0.5% triton and click chemistry was used to conjugate biotin-TEG-azide (Eurogentec) to EdU-labelled DNA. Cells were re-suspended in lysis buffer and sonication was performed using a Qsonica sonicator. Biotin conjugated DNA-protein complexes were captured using streptavidin beads (Ademtech). Captured complexes were washed with lysis buffer and high salt. Proteins associated with nascent DNA were eluted under reducing conditions by boiling into SDS sample buffer for 30 min at 95 °C.

### DNA fibers labelling

DNA fibers labelling was performed as previously described (Lossaint et al., 2013; Ribeyre et al., 2016). Cells were labeled with 25μM IdU, washed with warm media and exposed to 50 μM CldU. Cells were lysed and DNA fibers were stretched onto glass slides. The DNA fibers were denatured with 2.5 M HCl for 1 hour, washed with PBS and blocked with 2% BSA in PBS-Tween for 60 minutes. IdU replication tracts were revealed with a mouse anti-BrdU/IdU antibody from BD Biosciences (347580) and CldU tracts with a rat anti-BrdU/CldU antibody from Eurobio (ABC117-7513). The following secondary antibodies were used: alexa fluor 488 anti-mouse antibody (Life A21241) and Cy3 anti-rat antibody (Jackson Immunoresearch 712-166-153). Replication tracts lengths were analyzed using ImageJ software. For statistical analysis we used a non-parametrical Mann-Whitney with Prism software.

### Immunofluorescence

Cells were plated on glass coverslips and fixed with 4% paraformaldehyde in PBS for 20 min at room temperature. When indicated cells were incubated with EdU (5-ethynyl-2’-deoxyuridine) for the indicated times. PFA-fixed cells were permeabilized with 0.2% Triton X-100 in PBS for 5 min. Primary (Ser139 Phospho-H2A.X ; Millipore 05-636 and RIF1 ; Bethyl A300-568A-M) and secondary antibodies (anti-mouse Alexa 488 and anti-rabbit alexa 546) were prepared in PBS with 0.1% Tween and incubations were carried out in a humidified chamber at room temperature (60 min and 30 min, respectively). EdU was coupled with Alexa fluor 555 using Click chemistry. DNA was stained with Hoechst. The cells were mounted on glass slides with Prolong (Life). Cells were analyzed by fluorescence microscopy and quantification of various signals was performed using CellProfiler software (Carpenter et al., 2006).

### Proximity Ligation Assay

Cells were plated on glass coverslips and fixed with 4% paraformaldehyde in PBS for 20 min at room temperature. When indicated cells were incubated with EdU (5-ethynyl-2’-deoxyuridine). PFA-fixed cells were permeabilized with 0.5% Triton X-100 in PBS for 20 min. EdU was coupled with Alexa fluor 555 or biotin-TEG-azide using Click chemistry. Primary antibodies against SMC1 (Bethyl A300-055A), SMC3 (Bethyl A300-060A), PCNA (Sigma-Aldrich P8825), Biotin (Bethyl A150-109A or Jackson Immunoresearch 200-002-211) were incubated overnight. Probes from Duolink In Situ PLA Probe Anti-Rabbit PLUS (DUO92002, Sigma-Aldrich) and Duolink In Situ PLA Probe Anti-Mouse MINUS (DUO92004, Sigma-Aldrich) we incubated with coverslip 60 min at 37°C. For ligation (30 min at 37°C) and amplification (100 min at 37°C), Duolink In Situ Detection Reagents Green (DUO92014, Sigma-Aldrich) was used. The cells were mounted on glass slides with Duolink In Situ Mounting Medium with DAPI (DUO82040, Sigma-Aldrich). Cells were analyzed by fluorescence microscopy and quantification of PLA signal was performed using CellProfiler software (Carpenter et al., 2006).

### Flow cytometry

Cells were labelled with EdU for 15 min then fixed in 80% ethanol. After permeabilization, EdU was coupled with Alexa fluor 488 using Click chemistry. DNA was stained using propidium iodide and analysis was performed on Miltenyi MACS quant device.

### Mass Spectrometry Analysis

Mass spectrometry was performed as indicated in (Kumbhar et al., 2018). Analysis of raw files was performed using MaxQuant (Cox and Mann, 2008) version 1.5.6.5 using default settings with label-free quantification option enabled. Raw file spectra were searched against the human UniProt reference database. Protein, peptide, and site false discovery rate (FDR) were adjusted to < 0.01.

## Supplementary figures legends

**Sup Figure 1:**
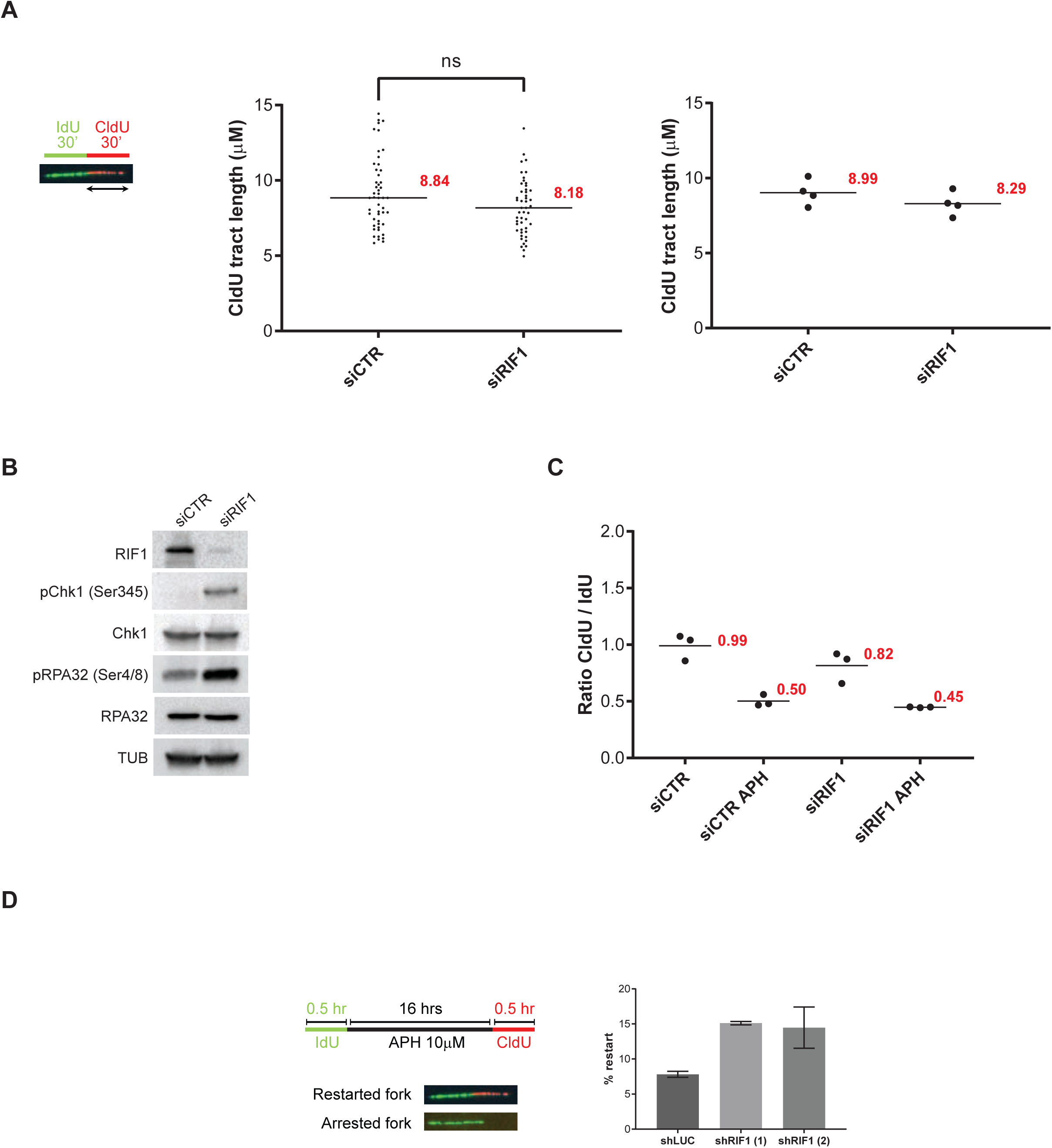
**A**. DNA fibers labelling. HeLa S3 cells were labelled for 30 min with IdU and then for 30 min with CldU. Graphic representation of the ratios of CldU tract length. The horizontal bar represents the median with the value indicated in red. For statistical analysis Mann-Whitney test was used; ns, non-significant. At least 50 replication tracts were measured for each experimental condition. The second graphic representation is showing the average of four independent experiments. **B**. Western-blot analysis of the indicated proteins upon transfection with siRNA directed against RIF1 or a control target. **C**. This graphic representation is showing the average of the three independent experiments from Figure 1D. **D**. Analysis of replication restart upon APH treatment using DNA fibers labelling. HeLa S3 cells were labelled for 30 min with IdU, then treated 16 hrs with 10μM APH and then for 30 min with CldU. Graphic representation of the percentage of restart based on 3 independent experiments.

**Sup Figure 2:**
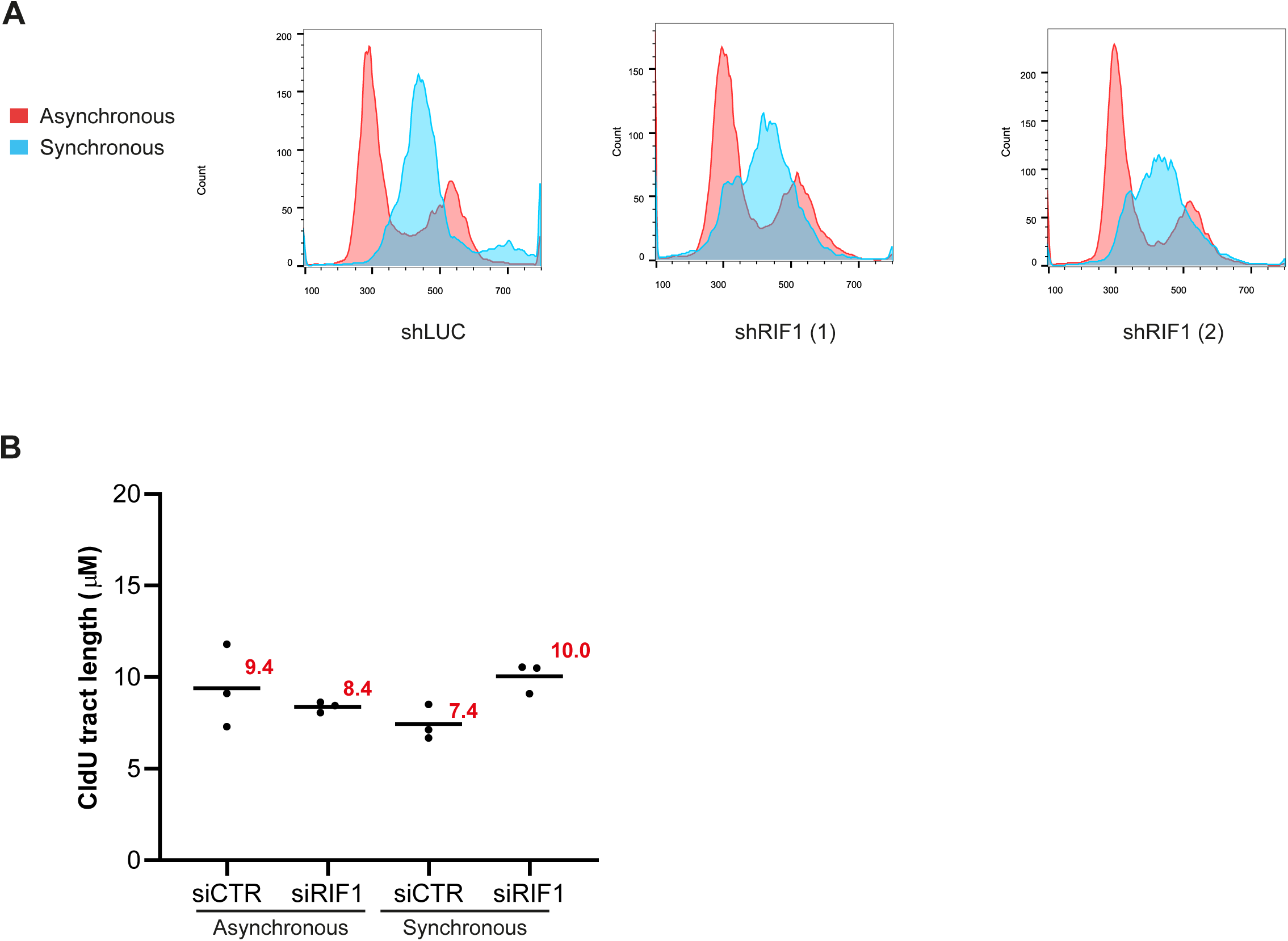
**A**. Flow cytometry analysis of cells depleted or not for RIF1 in asynchronous or synchronous conditions (18 hours thymidine block followed by 2 hours release). DNA was stained using propidium iodide. **B**. This graphic representation is showing the average of the three independent experiments from Figure 2F.

**Sup Figure 3:**
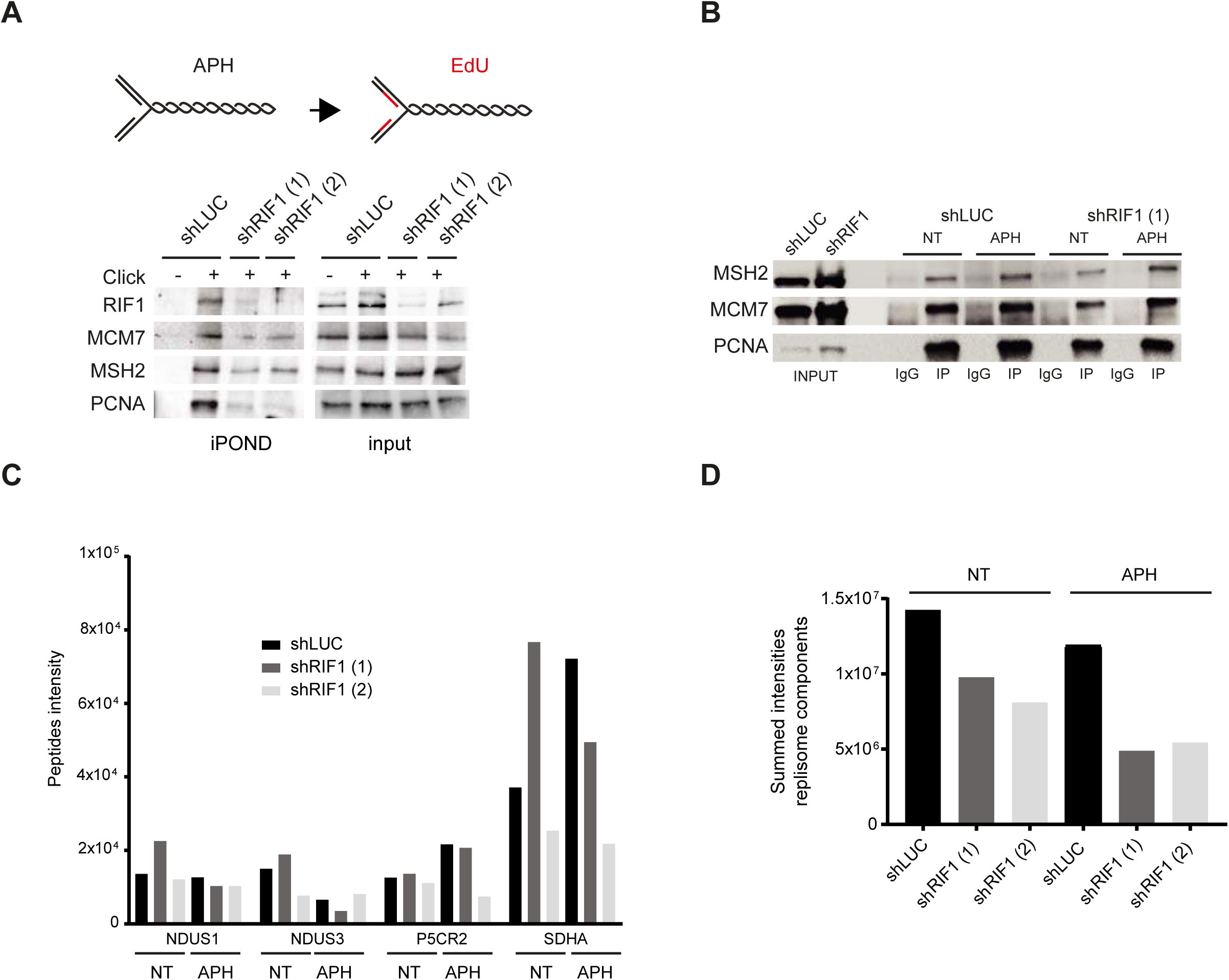
**A**. iPOND experiment. HeLa S3 cells (with shLUC or two different shRIF1) were treated 30 min with 0.1 μM aphidicolin (APH) then washed and labelled with EdU for 30 min. Indicated proteins were analyzed by Western-blotting. In no click samples, biotin-TEG azide was replaced by DMSO. **B**. Western-blot analysis of indicated proteins after immunoprecipitation with an antibody directed against PCNA or against mouse IgG. When indicated HeLa S3 cells (shLUc or shRIF1) were treated for 30 min with 0.1 μM aphidicolin (APH). **C**. Peptides intensity of proteins not specific from the replisome from the iPOND experiment from Figure 3C. **D**. Summed intensities of peptides corresponding to replisomes components (listed in Supplementary Table 2)from the iPOND experiment from Figure 3C.

**Sup Figure 4:**
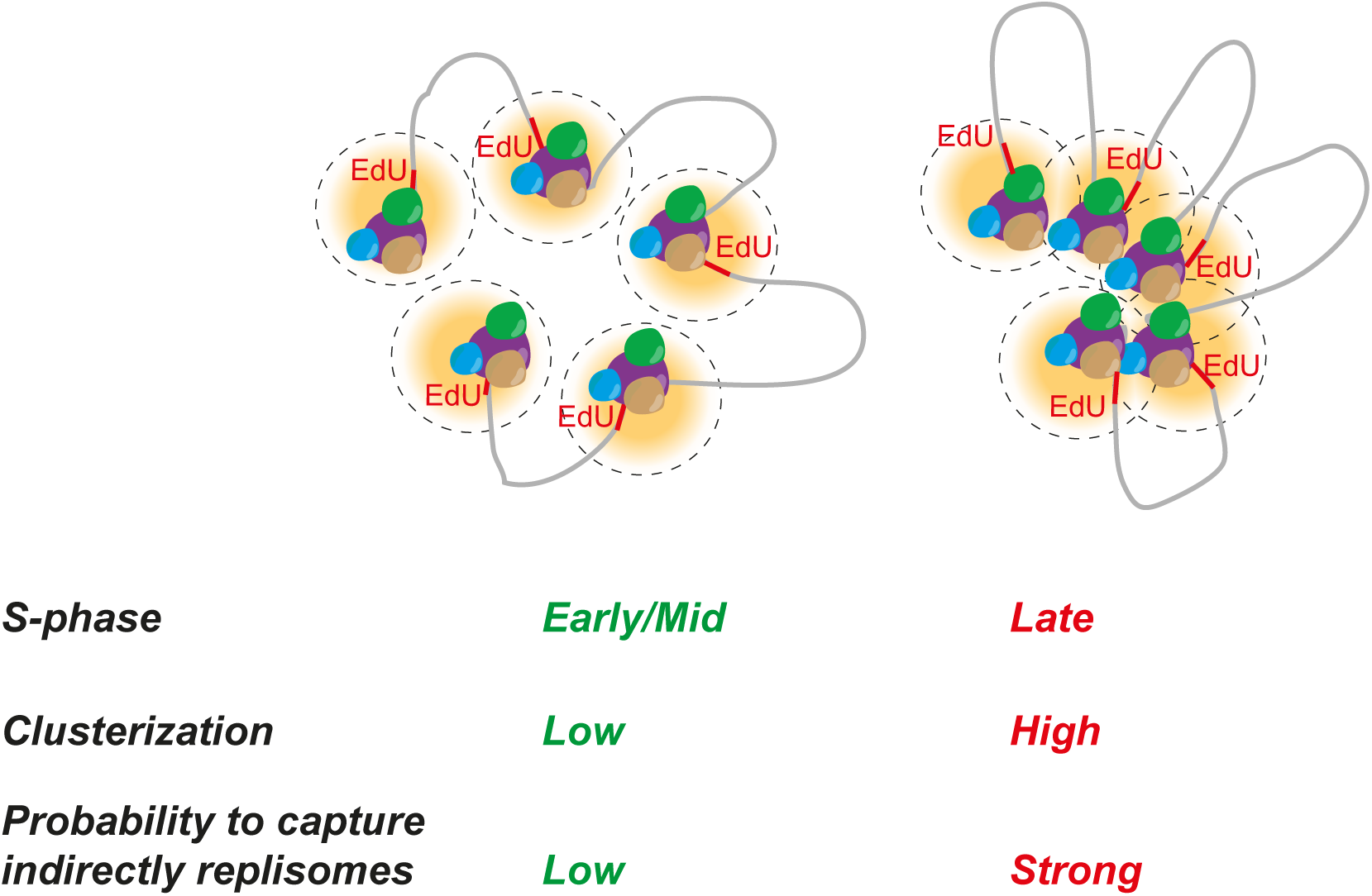
Scheme explaining how replication organization is impacting iPOND efficiency.

**Sup Figure 5:**
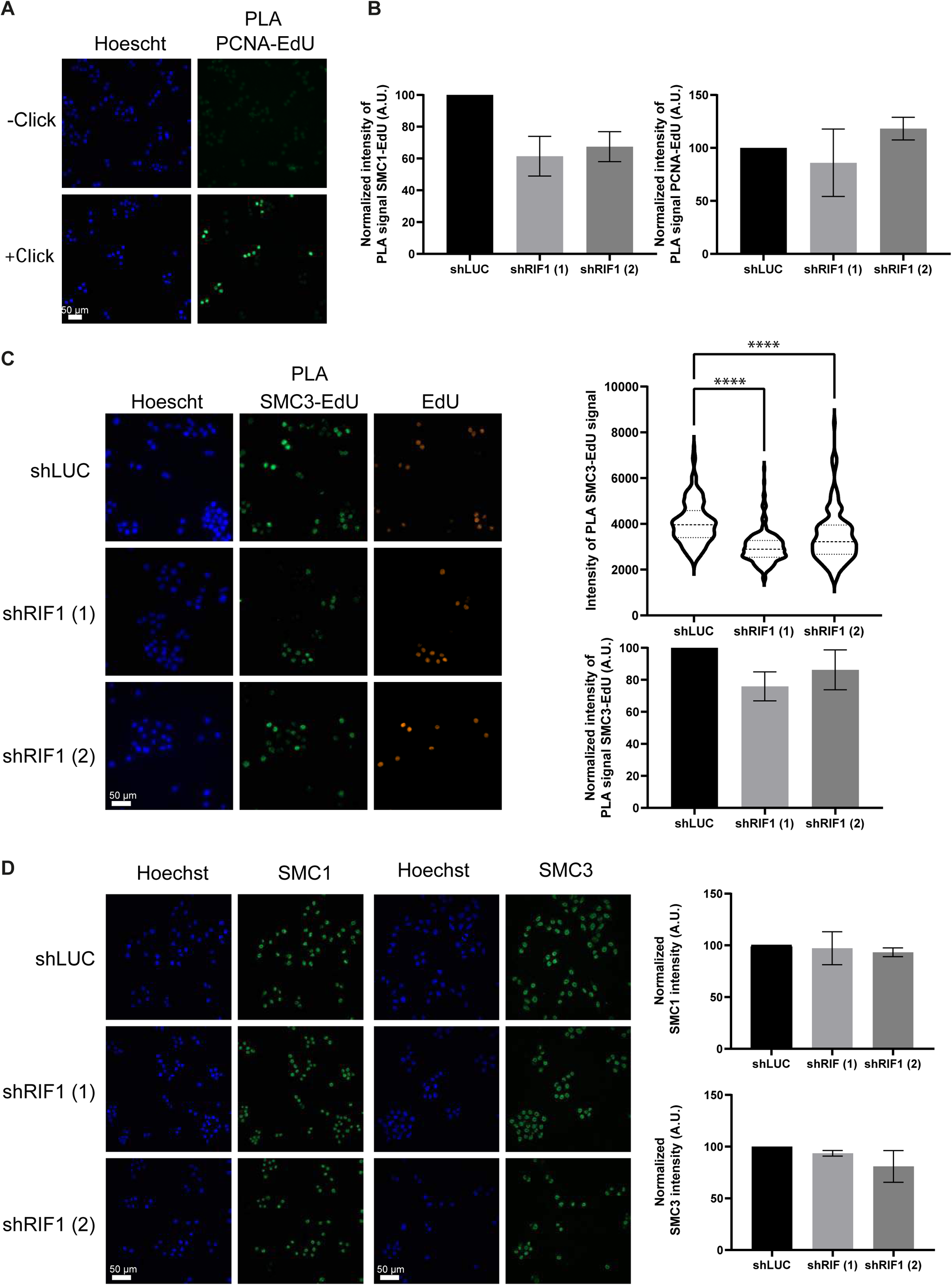
**A**. Immunofluorescence analysis of PLA signal between EdU and PCNA, in -click control the Biotin-TEG azide was replaced by DMSO. **B**. Graphic representation of the average PLA signal (normalized to 100 in shLUC) from 3 independent experiments corresponding to Figure 5C. **C**. Immunofluorescence analysis of PLA signal between EdU and SMC3 upon 30 min treatment with 0.1 μM APH in HeLa S3 cells expressing shRNAs against luciferase or RIF1. EdU-positive cells were labelled with Alexa-Fluor 555. The level of PLA signal within the nucleus was quantified using CellProfiler. Graphical representation of the PLA signal, at least 100 cells were quantified in each condition. For statistical analysis Mann-Whitney test was used; ****p<0.0001. Graphic representation of the average PLA signal (normalized to 100 in shLUC) from 3 independent experiments. **A**. Immunofluorescence analysis of SMC1 and SMC3. Graphic representation of the average SMC1 and SMC3 intensity within nucleus (normalized to 100 in shLUC) from 3 independent experiments.

